# An Interpretable Machine Learning Framework for Early- versus Late-Stage Classification of Breast Cancer from Gene Expression, with Co-expression Network Analysis

**DOI:** 10.64898/2026.10.18.759962

**Authors:** Sriporna Biswas, Tanjidul Huda

## Abstract

Anatomical stage is a principal determinant of treatment and prognosis in breast cancer, yet it is unclear how much information about stage is carried by the transcriptome of the primary tumour. We asked whether an a priori panel of 65 cancer-related genes can distinguish early-stage (AJCC Stage I–II) from late-stage (Stage III–IV) breast cancer, and which genes contribute to any signal. RNA-seq expression and pathologic stage for The Cancer Genome Atlas breast invasive carcinoma cohort (TCGA-BRCA) were obtained from the UCSC Xena platform. After restricting the data to primary tumours with a known stage, 1,075 tumours were analysed (805 early, 270 late) and split into stratified training (n = 860) and held-out test (n = 215) sets. Logistic regression, a support vector machine, random forest and gradient boosting were tuned by stratified five-fold cross-validation, with median imputation, standardisation and SMOTE oversampling fitted on training data only and classification thresholds chosen on out-of-fold training predictions. On the test set, Logistic Regression had the highest ROC-AUC, 0.618 (95% CI 0.530–0.704), with balanced accuracy 0.578, sensitivity 0.741 and specificity 0.416; differences in AUC between models were not statistically significant (DeLong p = 0.199–0.585). Permutation importance in the Gradient Boosting model ranked *POU5F1*, *FLT3* and *NOTCH1* highest, but this model’s AUC confidence interval included 0.5, so the ranking is exploratory. No gene was differentially expressed between stage groups after false-discovery-rate (FDR) correction (smallest FDR = 0.057, *CD44*). A co-expression network built from training tumours identified *REL*, *EP300* and *ESR1* as the most connected genes, a property that reflects co-expression rather than regulation or association with stage. Limitations include a single cohort, no external validation, no clinical covariates and a small number of late-stage test tumours. Expression of this gene panel carries limited but measurable information about anatomical stage; expression-based stage models should be combined with clinicopathological variables and validated in independent cohorts before any clinical application is considered.

## INTRODUCTION

Breast cancer is the most commonly diagnosed cancer in women worldwide, with approximately 2.3 million new cases and 666,000 deaths in 2022 [1]. Its anatomical stage, which the American Joint Committee on Cancer (AJCC) defines from tumour size, regional lymph-node involvement and distant metastasis, is central to treatment planning and prognosis [2]. Patients with locally advanced (Stage III) or metastatic (Stage IV) disease generally require more intensive therapy and have poorer survival than those diagnosed at Stage I–II.

Large genomic projects such as The Cancer Genome Atlas (TCGA) have shown that breast tumours are molecularly heterogeneous [3], and expression-based signatures such as PAM50 stratify patients by intrinsic subtype and risk of relapse [4]. Machine learning (ML) is now widely used to classify tumours from such data [5–7], and recent studies integrate multi-omics data with graph neural networks for cancer-type and breast cancer subtype classification [8, 9]. Three related prediction tasks should be distinguished. Molecular subtype classification assigns tumours to classes that are themselves defined by gene expression; survival or relapse prediction models a time-to-event outcome that also depends on treatment and follow-up; and anatomical stage classification predicts a clinicopathological label defined by tumour size, nodal status and metastasis, independently of the transcriptome [2]. Whether, and to what degree, the expression profile of the primary tumour reflects its anatomical extent is therefore a distinct question. A strong association would point to expression programmes that accompany tumour progression, whereas a weak association would indicate that stage carries information that bulk expression does not readily capture. Network-based studies suggest that expression changes between breast cancer conditions are often subtle and distributed across interacting genes [10], which makes a careful, quantitative answer to this question worthwhile.

In this study we developed an interpretable, leakage-aware ML framework to distinguish early-stage (I–II) from late-stage (III–IV) breast cancer using 65 a priori selected cancer-related genes in TCGA-BRCA. The study makes four contributions: (i) an explicitly defined cohort of primary tumours with a reproducible staging rule; (ii) an evaluation pipeline in which imputation, scaling and SMOTE are fitted on training data only, decision thresholds are tuned on out-of-fold predictions, and performance is reported with confidence intervals and imbalance-robust metrics; (iii) a comparison of four classifiers with statistical testing of AUC differences; and (iv) a side-by-side examination of three complementary views of the same genes—model-based permutation importance, univariate differential expression and co-expression network centrality. Rather than proposing new biomarkers, the aim is to quantify how much stage information this panel carries and to describe, with appropriate caution, which genes contribute to that signal.

## BACKGROUND

Gene expression profiling measures the activity of thousands of genes in a tumour and therefore captures molecular processes that are not visible in histology [3, 6]. Such data pose specific challenges for ML: the number of features is large relative to the number of samples, which raises the risk of over-fitting; measurements are noisy; many genes are strongly correlated; and clinically important classes are often under-represented [5]. Classical models address these problems in different ways. Regularised logistic regression provides an interpretable linear baseline, support vector machines with kernels capture non-linear decision boundaries [11], and tree ensembles such as random forests [12] and gradient boosting [13] model interactions between features.

Four methodological issues are particularly relevant to stage classification. First, stage classes are imbalanced, so accuracy is dominated by the majority class and can appear high even when a model predicts a single class [14, 15]. Second, oversampling with the synthetic minority over-sampling technique (SMOTE) [16], or scaling fitted before the train–test split, leaks information from test samples into training and inflates performance [17, 18]. Third, flexible models are difficult to interpret, which limits biological insight; model-agnostic approaches such as permutation importance [12] and SHapley Additive exPlanations (SHAP) [19, 20] attribute predictive performance or individual predictions to features, and interpretable models have been used to identify expression biomarkers in early-stage disease [21]. Fourth, predictive importance, statistical association and network centrality answer different questions: a gene can improve prediction in combination with other genes without being individually differentially expressed, and a highly connected gene in a correlation network need not be associated with the outcome at all. Treating these three properties as interchangeable evidence of biological relevance can therefore be misleading.

## LITERATURE REVIEW

Kourou et al. reviewed ML applications in cancer prognosis and prediction and emphasised the risk of over-fitting when samples are few and features many [5]. Subsequent work has achieved high accuracy on tasks in which classes differ strongly in expression. Ensemble learning combined with feature selection has been applied to cancer-type classification from gene expression [7], and graph neural networks that integrate several omics layers have improved multi-omics cancer classification [8] and breast cancer subtyping [9]. These studies separate tissue types or molecular subtypes, whose definitions are closely tied to expression; their performance therefore does not indicate how well expression can predict anatomical stage within a single cancer type, a label that reflects anatomical spread rather than a distinct molecular programme.

A second line of work focuses on identifying genes and networks associated with disease state. Wang and Liu identified breast cancer biomarkers through rewiring of gene regulatory networks between conditions [10], and ECMarker used an interpretable model to identify gene expression biomarkers that predict clinical outcomes in early disease stages [21]. Gene set enrichment analysis [22] and KEGG pathway resources [23] are widely used to place candidate genes in a functional context; pathway enrichment was not performed in the present study. A third, methodological line of work concerns evaluation under class imbalance. Blagus and Lusa showed that SMOTE offers limited benefit for high-dimensional data unless it is combined with variable selection [18], and guidance on imbalanced evaluation recommends precision–recall analysis and the Matthews correlation coefficient over accuracy [14, 15].

The present study differs from these works in its question and design. It addresses anatomical stage rather than tissue type, molecular subtype or outcome; it uses a single public cohort and a fixed gene panel; and it combines leakage-aware evaluation, imbalance-robust metrics, model explanation, univariate differential expression and co-expression network analysis within one framework, while keeping these lines of evidence explicitly separate. We do not claim that this combination is unique; rather, it provides a transparent and reproducible baseline against which richer models and data can be compared.

## METHODOLOGY

### Study Design and Workflow

Figure 1 summarises the workflow. The cohort was defined first and split once into training and test sets. All steps that learn from data for prediction—imputation, scaling, oversampling, hyper-parameter tuning and threshold selection—used the training set only, and the test set was used once for final evaluation. Two descriptive analyses were run alongside the predictive analysis: differential expression on all analysed tumours and a co-expression network on the training tumours only (details below).

**Figure 1:**
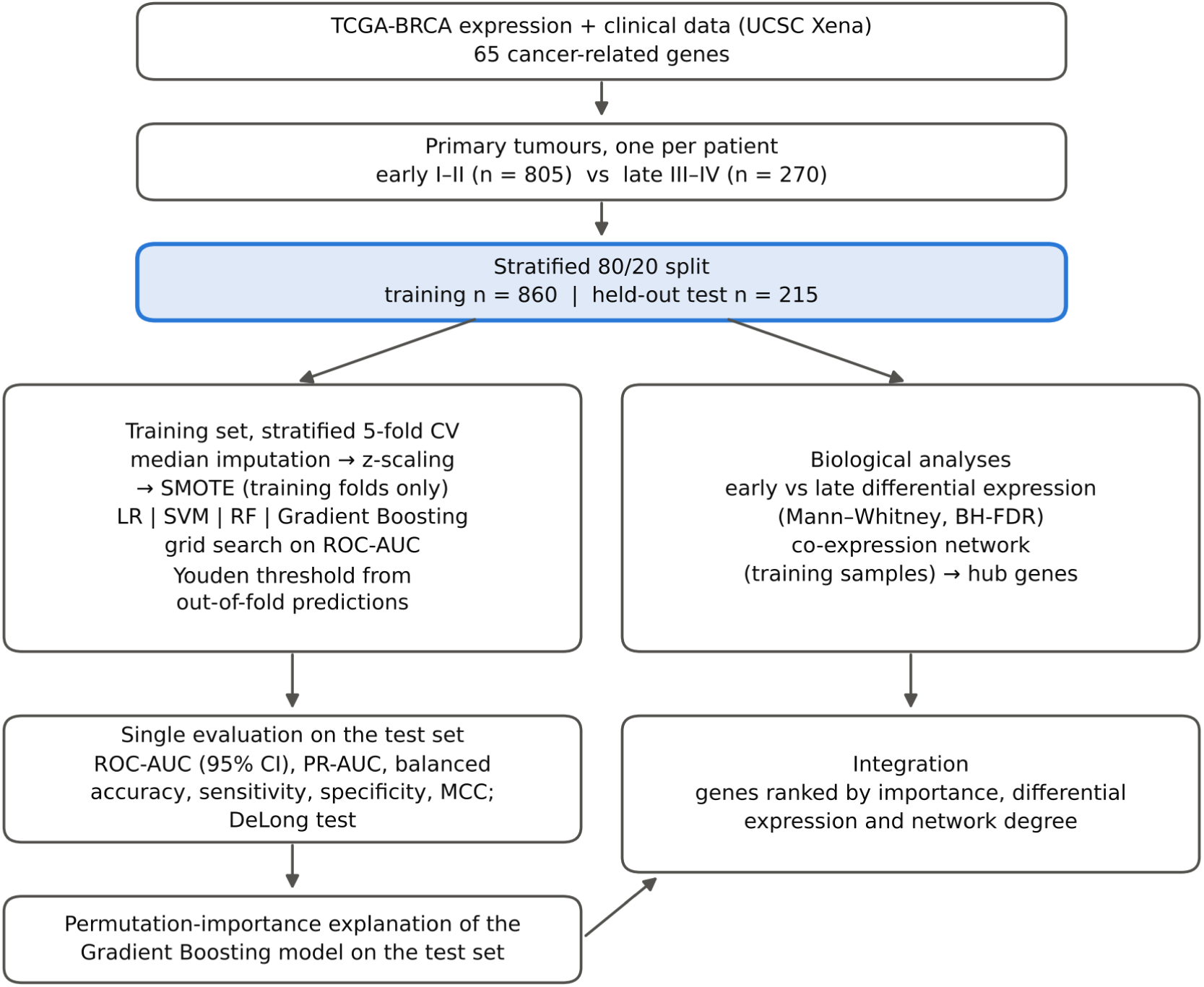
Study workflow. Imputation, scaling, SMOTE, hyper-parameter tuning and threshold selection used training data only, and the held-out test set was used once for final evaluation. As described in the Methods, differential expression was computed on all 1,075 analysed tumours and the co-expression network on the 860 training tumours only. The “Integration” box denotes a qualitative comparison of the resulting gene lists, not a formal statistical integration.

### Data Source

Gene expression and clinical data for the TCGA breast invasive carcinoma cohort (TCGA-BRCA) [3] were downloaded from the UCSC Xena browser [24] as a single sample-by-feature table for 65 genes. Expression values were taken from the TCGA hub dataset TCGA.BRCA.sampleMap/HiSeqV2_PANCAN (IlluminaHiSeq pan-cancer normalised RNA-seq; RSEM normalised counts transformed as log2(x + 1) and mean-centred per gene across all TCGA cohorts; version 13 October 2017), and AJCC pathologic stage was taken from the TCGA-BRCA phenotype matrix (TCGA.BRCA.sampleMap/BRCA_clinicalMatrix; version 6 December 2019). The genes were chosen a priori from cancer-related signalling, cell-cycle, DNA-repair, transcriptional-regulation and stem-cell pathways (Supplementary Table S1). All 65 genes were used as model inputs; no data-driven feature selection was performed. The data are public and de-identified, so no ethical approval was required.

### Cohort and Outcome Definition

The downloaded table contained 1,247 samples. Primary solid-tumour samples (TCGA barcode sample type 01; n = 1,101) were retained, and 139 solid normal tissue samples and 7 metastatic samples were removed, because normal or metastatic tissue carries the patient’s stage label but not a primary-tumour expression profile. No patient contributed more than one primary-tumour sample (n = 1,101). Tumours were labelled early stage (AJCC Stage I, IA, IB, II, IIA or IIB; class 0) or late stage (Stage III, IIIA, IIIB, IIIC or IV; class 1) [2]. Tumours with unknown (Stage X), discrepant or missing stage were excluded (n = 26: Stage X, 14; missing, 9; discrepant, 3), leaving 1,075 tumours. No gene had more than 20% missing values; the remaining missing values (0.19% of cells) were imputed with the training-set median inside the modelling pipeline.

### Data Splitting, Pre-processing and Class Balancing

The cohort was split once into a stratified training set (80%, n = 860) and a held-out test set (20%, n = 215) with a fixed random seed, preserving the class ratio in both sets. The test set was not used for imputation, scaling, oversampling, tuning or threshold selection. Pre-processing was implemented as a pipeline in which resampling acts only during fitting, as in imbalanced-learn [25], and was re-fitted within each cross-validation fold: median imputation, z-score standardisation, and SMOTE (k = 5 nearest neighbours) to oversample late-stage tumours to parity with early-stage tumours [16]. SMOTE was therefore applied only to the data used for fitting; validation folds and the test set kept their natural class ratio. After tuning, the selected configuration of each model was refitted on the full training set through the same pipeline, so that the final models were trained on 1,288 samples (644 per class) after oversampling. The prespecified primary analysis used SMOTE. As a sensitivity analysis, every model was also trained without SMOTE, using class weights inversely proportional to class frequency, the same hyper-parameter grids and the same threshold-selection procedure.

### Classification Models and Hyper-parameter Tuning

Four classifiers were compared: L2-regularised logistic regression (LR) as a linear baseline [26]; a support vector machine with a radial-basis-function kernel (SVM) [11], with probability estimates enabled; random forest (RF) with 500 trees [12]; and gradient-boosted decision trees [13] (scikit-learn HistGradientBoostingClassifier, without early stopping). Hyper-parameters were selected by grid search with shuffled, stratified five-fold cross-validation on the training set, maximising ROC-AUC (grids and selected values in Supplementary Table S2). The random seed was fixed at 42 for data splitting, cross-validation, SMOTE and all stochastic models.

### Decision Threshold and Performance Evaluation

Because the default probability threshold of 0.5 is unsuitable for imbalanced data, each model’s threshold was set to maximise Youden’s J (sensitivity + specificity − 1) [27] on out-of-fold predictions for the training set, obtained by five-fold cross-validation of the selected configuration, and was then applied unchanged to the test set. Late stage was treated as the positive class. Test-set performance was summarised by ROC-AUC, precision–recall AUC (average precision) [14], balanced accuracy, sensitivity (recall for late stage), specificity, precision, F1-score and the Matthews correlation coefficient (MCC) [15]. Ninety-five per cent confidence intervals (CIs) for ROC-AUC and balanced accuracy were obtained by the percentile method from 1,000 bootstrap resamples of the test set; these intervals reflect test-set sampling variability only. The AUC of the model with the highest test AUC was compared with that of each other model using DeLong’s two-sided test for correlated ROC curves [28], without adjustment for the three comparisons. Cross-validated AUC (mean ± SD across the five folds) is reported for the selected configuration of each model; because it was computed on the same folds used for tuning, without nested cross-validation, it may be slightly optimistic and is reported as a stability check rather than an unbiased estimate.

### Model Interpretability

Feature importance was estimated for the tuned Gradient Boosting model by model-agnostic permutation importance [12] on the test set: each gene was randomly permuted 30 times and the mean decrease in ROC-AUC was recorded, with negative means set to zero. The direction of effect was summarised by the Spearman correlation between a gene’s expression and the model’s predicted probability of late stage. Permutation importance describes how the model uses each gene for prediction; it is not a test of association between the gene and stage. The analysis was not repeated for the other three classifiers, including Logistic Regression.

### Differential Expression Analysis

Expression of each gene was compared between early-stage (n = 805) and late-stage (n = 270) tumours in the full analysed cohort (n = 1,075) with the two-sided Mann–Whitney U test. Effect size was expressed as the rank-biserial correlation r (positive when expression is higher in late stage) and as the difference in mean log2 expression (late minus early). P-values were adjusted across the 65 genes with the Benjamini–Hochberg procedure [29], and genes with a false discovery rate (FDR) below 0.05 were considered stage-associated. This descriptive analysis was not used for model training or gene selection, so including test-set tumours does not affect the evaluation of the classifiers.

### Gene Co-expression Network and Hub Gene Identification

Pairwise Pearson correlations between all 65 genes were computed on the 860 training tumours. Undirected co-expression networks were built in NetworkX [30] by connecting gene pairs with |r| ≥ 0.7 (strong), |r| ≥ 0.5 (moderate) and |r| ≥ 0.3 (weak). Hub genes were ranked by degree (number of connections) in the network at |r| ≥ 0.5, with a prespecified fallback to |r| ≥ 0.3 if the 0.5 network had fewer than ten edges; the fallback was not needed. For Figure 7, genes were ordered by average-linkage hierarchical clustering of the correlation matrix. Correlation networks describe co-expression and do not establish regulatory relationships or their direction [31].

### Software and Reproducibility

Analyses were run in Python 3.13.16 with scikit-learn 1.9.1, pandas 3.0.5, SciPy and NetworkX 3.6.1; SMOTE was implemented as described by Chawla et al. [16]. The random seed was fixed at 42 throughout.

## RESULTS AND DISCUSSION

### Cohort Characteristics

After filtering, 1,075 primary tumours were analysed: 805 early-stage and 270 late-stage (25.1% late; Table 1, Figure 2). Stage IIA (n = 358) and Stage IIB (n = 258) were the most frequent early-stage categories, and Stage IIIA (n = 156) the most frequent late-stage category. The stratified split gave 860 training tumours (644 early, 216 late) and 215 test tumours (161 early, 54 late). Because of the imbalance, a trivial classifier that labels every tumour as early stage would reach 74.9% accuracy on the test set (161/215) while having no discriminative value; accuracy alone is therefore not informative here, and the analysis focuses on ROC-AUC, balanced accuracy, sensitivity, specificity and MCC.

**Figure 2:**
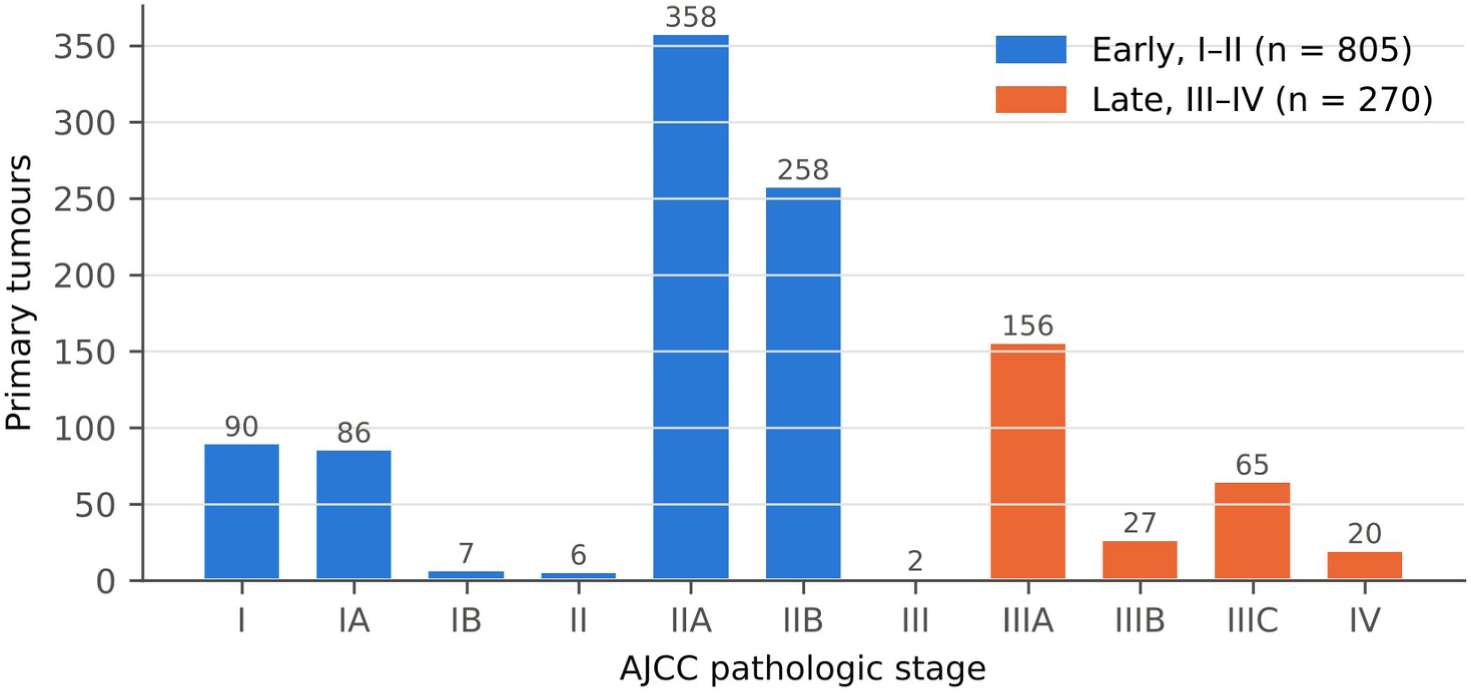
Distribution of AJCC pathologic stage among the 1,075 analysed primary tumours. Blue bars, early stage (I–II; n = 805); orange bars, late stage (III–IV; n = 270).

**Table 1:** Cohort characteristics and data partition. Percentages refer to the 1,075 analysed tumours. The last row gives the size of the training set after SMOTE oversampling.

| Characteristic | Value |
| --- | --- |
| Source | TCGA-BRCA via UCSC Xena |
| Samples in downloaded table | 1,247 |
| Primary tumours (sample type 01) | 1,101 |
| One sample per patient | 1,101 |
| Excluded: stage X, discrepant or missing | 26 |
| Analysed tumours | 1,075 |
| Early stage, I–II (class 0) | 805 (74.9%) |
| Late stage, III–IV (class 1) | 270 (25.1%) |
| Genes in panel | 65 |
| Training set (early / late) | 860 (644 / 216) |
| Test set (early / late) | 215 (161 / 54) |
| Training set after SMOTE (per class) | 1,288 (644) |

### Model Performance

Table 2 summarises test-set performance at each model’s tuned threshold. Logistic Regression had the highest ROC-AUC, 0.618 (95% CI 0.530–0.704), indicating modest discrimination between early- and late-stage tumours; the other models reached 0.589–0.595. The lower bound of the AUC confidence interval was above 0.5 for Logistic Regression (0.530) and, only marginally, for SVM (RBF) (0.506), whereas the intervals for Random Forest (0.499–0.678) and Gradient Boosting (0.498–0.689) included 0.5. DeLong tests comparing Logistic Regression with each other model gave p = 0.199 (SVM (RBF)), p = 0.497 (Random Forest) and p = 0.585 (Gradient Boosting), so the differences between models were not statistically significant. The ranking of models also differed between the test set and cross-validation, in which SVM (RBF) had the highest mean AUC (0.622 ± 0.025 vs 0.611 ± 0.036 for Logistic Regression). Logistic Regression is therefore described as the model with the highest test AUC rather than as a definitively superior model.

**Table 2:** Performance of the four classifiers on the held-out test set (n = 215; late stage = positive class). Threshold-dependent metrics were computed at each model’s Youden threshold selected on out-of-fold training predictions. PR-AUC, area under the precision– recall curve (average precision); Bal. acc., balanced accuracy; Sens., sensitivity; Spec., specificity; MCC, Matthews correlation coefficient. Values in parentheses are 95% bootstrap confidence intervals. CV AUC, five-fold cross-validated ROC-AUC on the training set (mean ± SD).

| Model | ROC-AUC (95% CI) | PR-AUC | Bal. acc. (95% CI) | Sens. | Spec. | F1 | MCC | CV AUC |
| --- | --- | --- | --- | --- | --- | --- | --- | --- |
| Logistic Regression | 0.618 (0.530–0.704) | 0.351 | 0.578 (0.512–0.648) | 0.741 | 0.416 | 0.426 | 0.140 | 0.611 ± 0.036 |
| SVM (RBF) | 0.595 (0.506–0.681) | 0.373 | 0.579 (0.503–0.658) | 0.556 | 0.602 | 0.405 | 0.138 | 0.622 ± 0.025 |
| Gradient Boosting | 0.594 (0.498–0.689) | 0.337 | 0.564 (0.490–0.640) | 0.537 | 0.590 | 0.389 | 0.111 | 0.582 ± 0.057 |
| Random Forest | 0.589 (0.499–0.678) | 0.385 | 0.528 (0.478–0.577) | 0.889 | 0.168 | 0.407 | 0.068 | 0.583 ± 0.062 |

At its tuned threshold of 0.36, Logistic Regression identified 40 of 54 late-stage test tumours (sensitivity 0.741) and correctly classified 67 of 161 early-stage tumours (specificity 0.416; Figure 4), with balanced accuracy 0.578 (95% CI 0.512–0.648) and MCC 0.140. The Youden threshold thus favoured sensitivity: 94 early-stage tumours were predicted as late stage, giving a precision of 0.299 (40/134) and an overall accuracy of 0.498 (107/215), both derived from the confusion matrix in Figure 4. Its precision–recall AUC was 0.351, against a no-skill baseline equal to the late-stage prevalence of 0.251 (Figure 3b). Cross-validated AUCs on the training set were within 0.03 of the test AUCs for all four models (Table 2), suggesting that the test-set estimates are not markedly unstable, within the caveats noted in the Methods.

**Figure 3:**
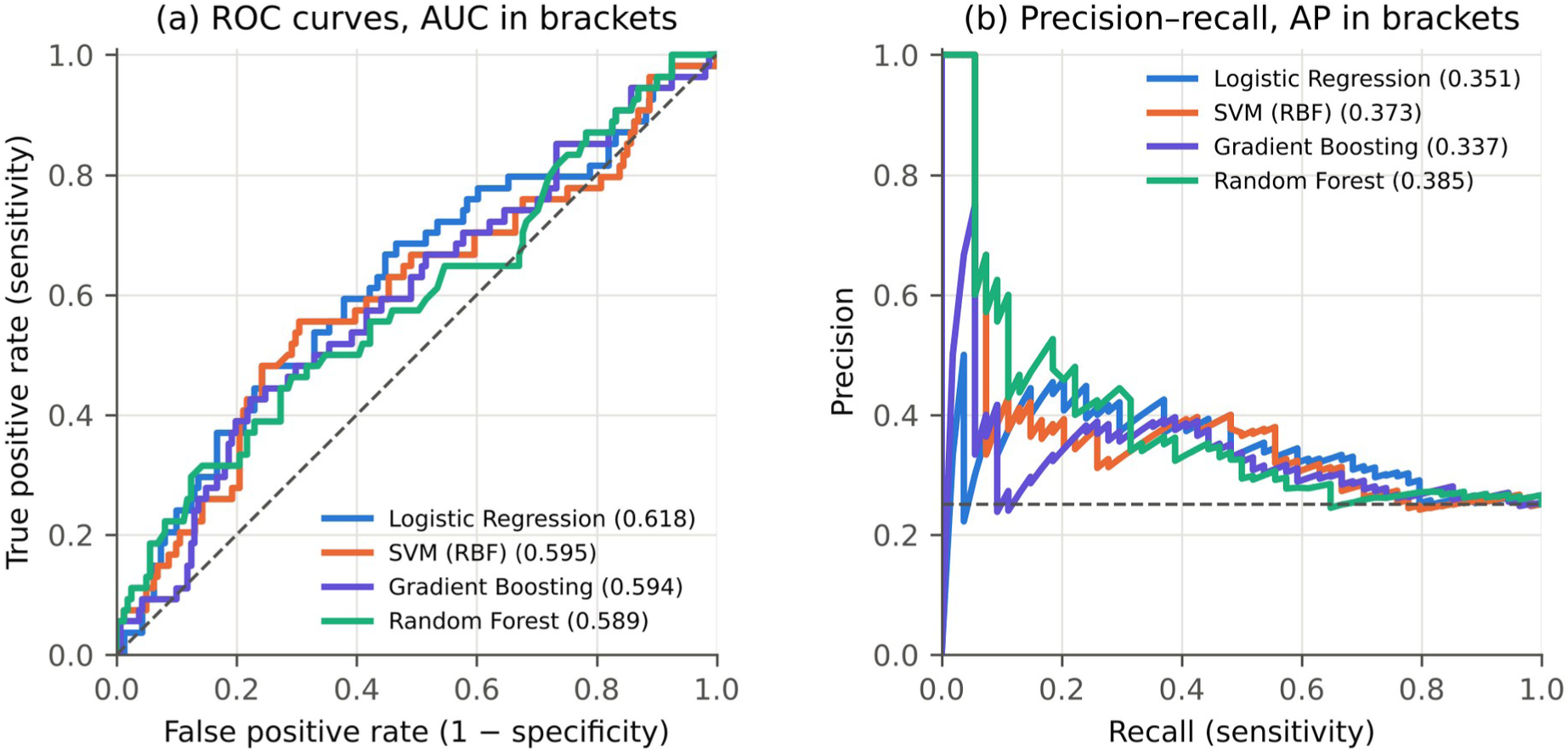
(a) Receiver operating characteristic (ROC) curves and (b) precision–recall curves of the four classifiers on the held-out test set (n = 215). Legend values are ROC-AUC in (a) and average precision (PR-AUC) in (b). Dashed lines indicate chance performance: the diagonal in (a) and the late-stage prevalence (0.251) in (b).

**Figure 4:**
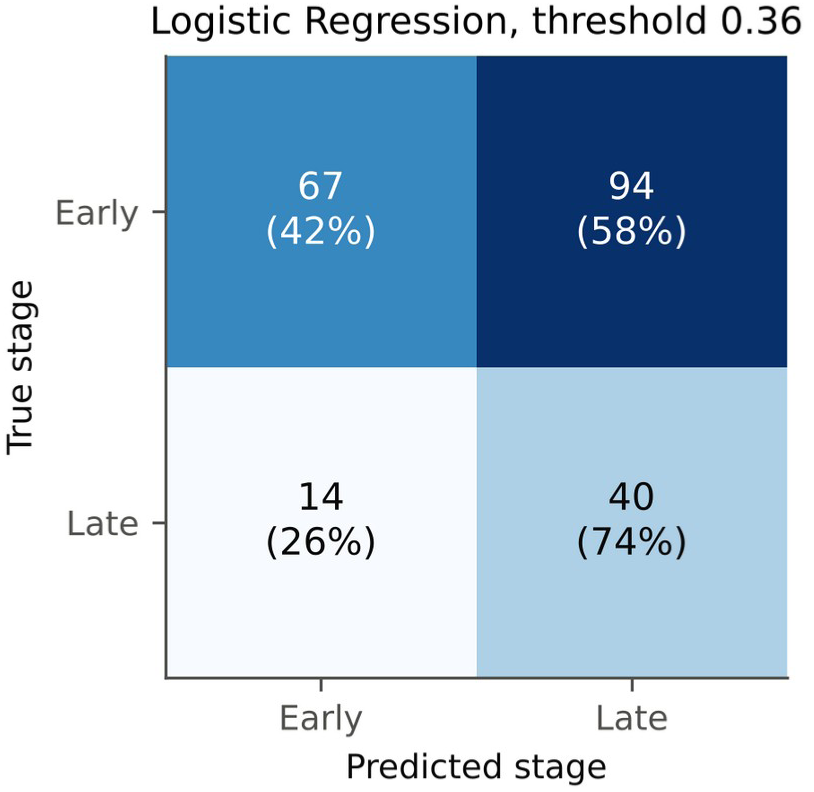
Confusion matrix of Logistic Regression on the held-out test set (n = 215) at the threshold (0.36) selected on out-of-fold training predictions. Percentages are calculated within each true class (row-wise).

### Sensitivity Analysis: Class Weighting Instead of SMOTE

The prespecified primary analysis used SMOTE within the training data, and all results in Table 2 and in the main text refer to that analysis. As a sensitivity analysis, each model was refitted with class weighting instead of SMOTE, using the same hyper-parameter grids and the same threshold-selection procedure (Supplementary Table S3). In the sensitivity analysis, test AUCs were 0.651 for Logistic Regression, 0.625 for SVM (RBF), 0.615 for Random Forest and 0.576 for Gradient Boosting, compared with 0.618, 0.595, 0.589 and 0.594, respectively, in the primary analysis. Class weighting therefore gave numerically higher AUCs for three of the four models (by approximately 0.03 for Logistic Regression and SVM (RBF), and by 0.026 for Random Forest) and a lower AUC for Gradient Boosting (by 0.018). Class-weighted Logistic Regression also had a higher balanced accuracy (0.638 vs 0.578) and MCC (0.239 vs 0.140).

Confidence intervals and DeLong tests were not computed for the sensitivity analysis, so these differences have not been formally tested and should not be interpreted as evidence that either imbalance-handling strategy is superior. Under both strategies all AUCs remained modest (0.576– 0.651), so the main conclusion—that this gene panel carries limited information about anatomical stage—does not depend on the choice of strategy. The sensitivity analysis also shows that SMOTE did not provide a consistent performance advantage over class weighting in this data set.

### Feature Importance in the Gradient Boosting Model

Figure 5 shows the genes with the largest permutation importance in the Gradient Boosting model (test AUC 0.594, 95% CI 0.498–0.689). The five most important genes were *POU5F1*, *FLT3*, *NOTCH1*, *FOS* and *NFKB2*, and higher expression of *POU5F1*, *FLT3*, *NOTCH1*, *FOS*, *NFKB2* and *PALB2* was associated with a lower predicted probability of late stage. All importance values were small: permuting any single gene reduced the test AUC by less than 0.05 on average (Figure 5). Two points limit the interpretation of this ranking. First, it was obtained from the Gradient Boosting model, whose AUC confidence interval included 0.5, rather than from Logistic Regression, the model with the highest test AUC. Second, permutation importance measures how much a fitted model relies on a gene, not whether that gene is associated with stage in the population. The ranking should therefore be regarded as exploratory and hypothesis-generating.

**Figure 5:**
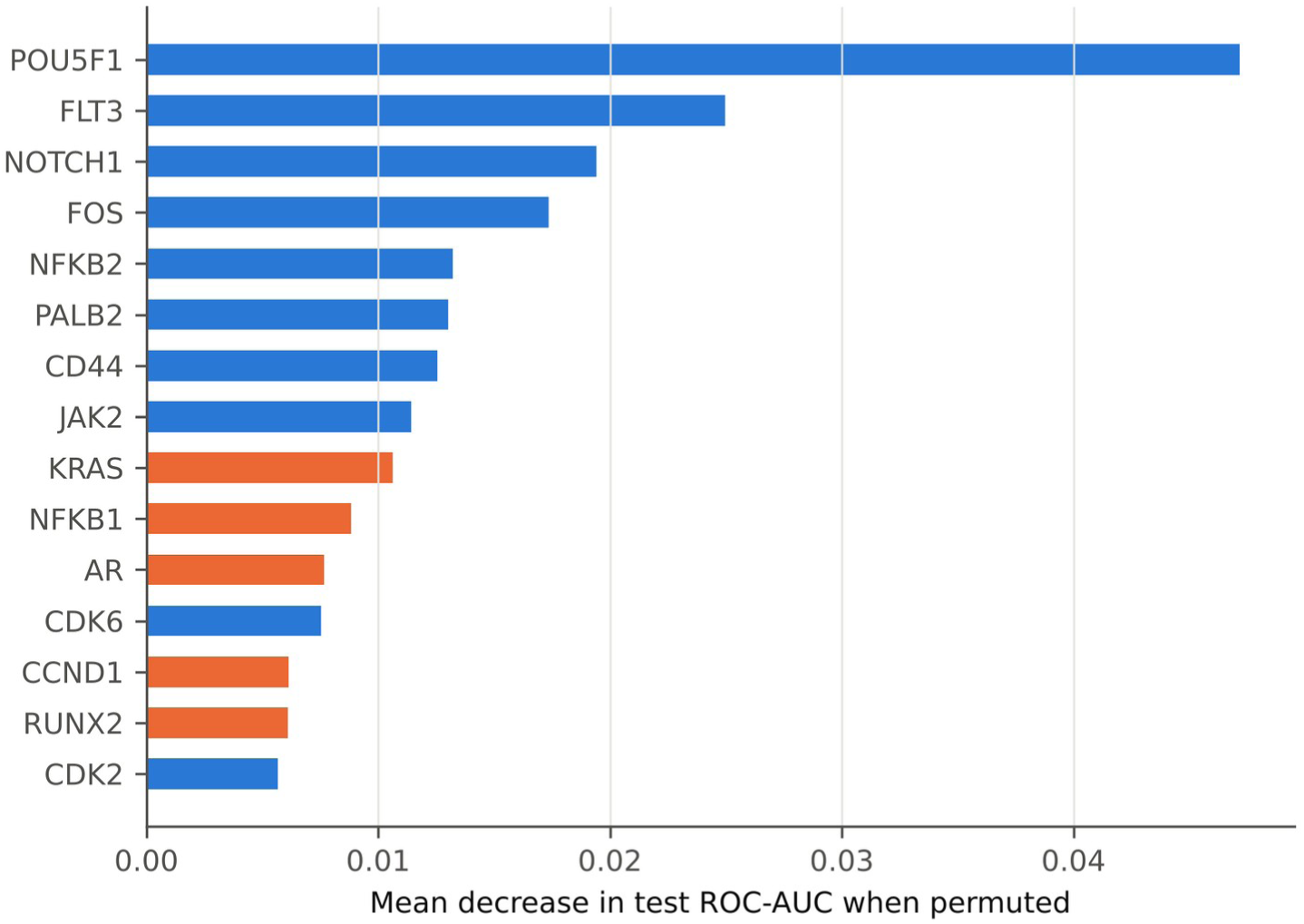
Permutation importance (mean decrease in test ROC-AUC over 30 permutations) of the 15 highest-ranked genes for the Gradient Boosting model on the held-out test set. Orange bars, higher expression associated with a higher predicted probability of late stage; blue bars, with early stage. Because this model’s AUC confidence interval included 0.5, the ranking is exploratory.

### Stage-associated Gene Expression

No gene reached the prespecified significance threshold of FDR < 0.05 for differential expression between early- and late-stage tumours (Table 3; full results in Supplementary Table S4). The strongest univariate signal was for *CD44*, whose expression was lower in late-stage tumours (difference in means −0.24; r = −0.14; nominal p < 0.001); after correction for 65 tests its FDR was 0.057, just above the threshold, and it is therefore not reported as stage-associated. Seventeen genes had nominal p < 0.05 (Supplementary Table S4), more than the approximately three that would be expected by chance if no gene were associated with stage (an approximate comparison, because the tests are correlated), but no individual difference survived FDR correction and all effect sizes were small (largest |r| = 0.14). Consistent with these small effects, the heatmap in Figure 6 shows no clear block structure separating the two stage groups. Together, these results indicate a weak and diffuse association between the expression of these genes and stage, rather than a few strongly stage-specific genes.

**Figure 6:**
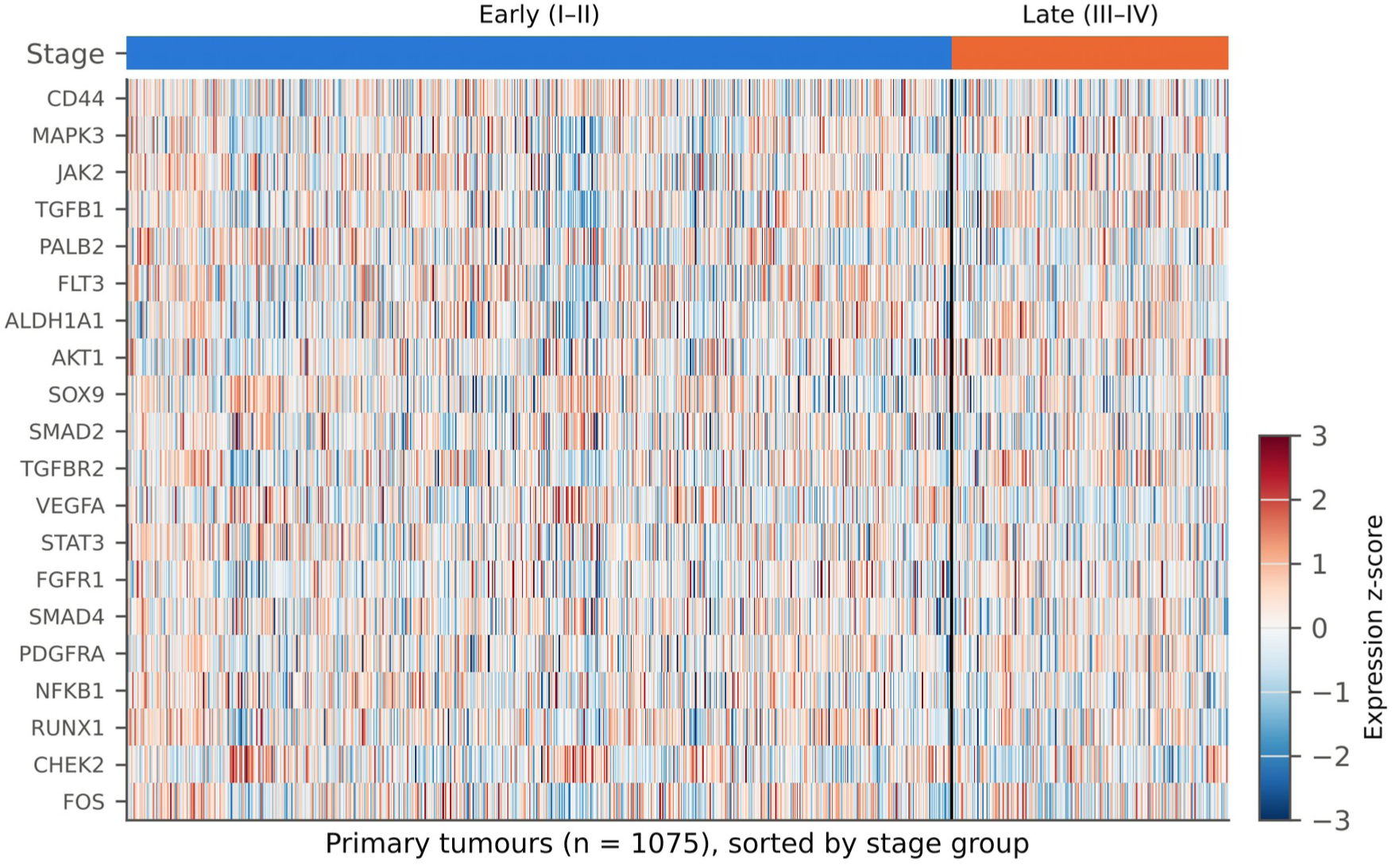
Standardised expression (z-score) of the 20 genes with the smallest differential-expression p-values across all 1,075 analysed tumours, ordered by stage group. The vertical line separates early-from late-stage tumours.

**Table 3:** Ten genes with the smallest p-values for differential expression between early- and late-stage tumours (Mann–Whitney U test, n = 1,075). Means are on the log2 expression scale; Difference, mean late minus mean early; r, rank-biserial correlation (positive, higher in late stage); FDR, Benjamini–Hochberg adjusted p-value across 65 genes. No gene reached FDR < 0.05.

| Gene | Mean early | Mean late | Difference | $r$ | $p$ | FDR |
| --- | --- | --- | --- | --- | --- | --- |
| *CD44* | 0.52 | 0.28 | −0.24 | −0.14 | < 0.001 | 0.057 |
| *MAPK3* | 0.07 | 0.20 | +0.13 | +0.12 | 0.005 | 0.10 |
| *JAK2* | 0.37 | 0.21 | −0.16 | −0.11 | 0.005 | 0.10 |
| *TGFB1* | −0.40 | −0.27 | +0.13 | +0.11 | 0.007 | 0.11 |
| *PALB2* | 0.36 | 0.27 | −0.09 | −0.10 | 0.010 | 0.11 |
| *FLT3* | 2.13 | 1.74 | −0.38 | −0.10 | 0.014 | 0.11 |
| *ALDH1A1* | −1.89 | −1.65 | +0.24 | +0.10 | 0.014 | 0.11 |
| *AKT1* | 0.18 | 0.28 | +0.10 | +0.10 | 0.015 | 0.11 |
| *SOX9* | 0.73 | 0.54 | −0.20 | −0.10 | 0.017 | 0.11 |
| *SMAD2* | 0.23 | 0.15 | −0.08 | −0.10 | 0.017 | 0.11 |

### Gene Co-expression Network and Hub Genes

Correlations among the panel genes in the training set are shown in Figure 7. At |r| ≥ 0.7 the network contained 7 edges among 9 genes. Lowering the threshold to 0.5 and 0.3 gave 70 edges (36 genes) and 385 edges (58 genes), respectively. Hub genes were ranked in the |r| ≥ 0.5 network (Figure 8): the most connected were *REL* (degree 11), *EP300* (degree 10), *ESR1* (degree 9), *GATA3* (degree 8) and *PIK3CA* (degree 7). Several genes shared the same degree below these five (Figure 8b), so ranks beyond the top five are not unique. The five highest-degree genes did not overlap with the ten genes most important to the Gradient Boosting model, and none of them reached FDR < 0.05 in the differential-expression analysis. Network centrality therefore captures a property—how strongly a gene co-varies with other panel genes—that is distinct from both predictive importance and association with stage.

**Figure 7:**
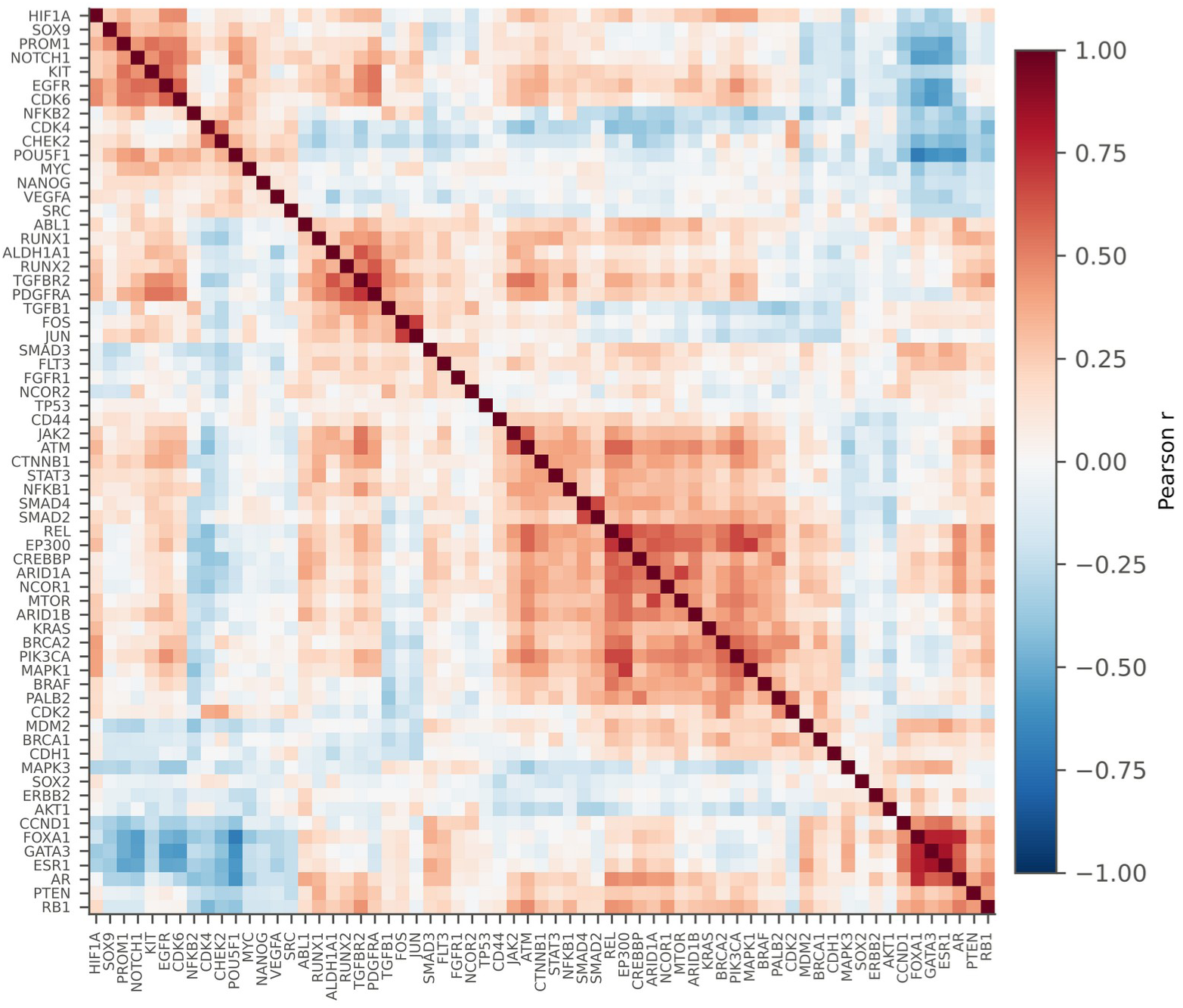
Pearson correlation matrix of the 65 panel genes computed on the 860 training tumours, with genes ordered by average-linkage hierarchical clustering.

**Figure 8:**
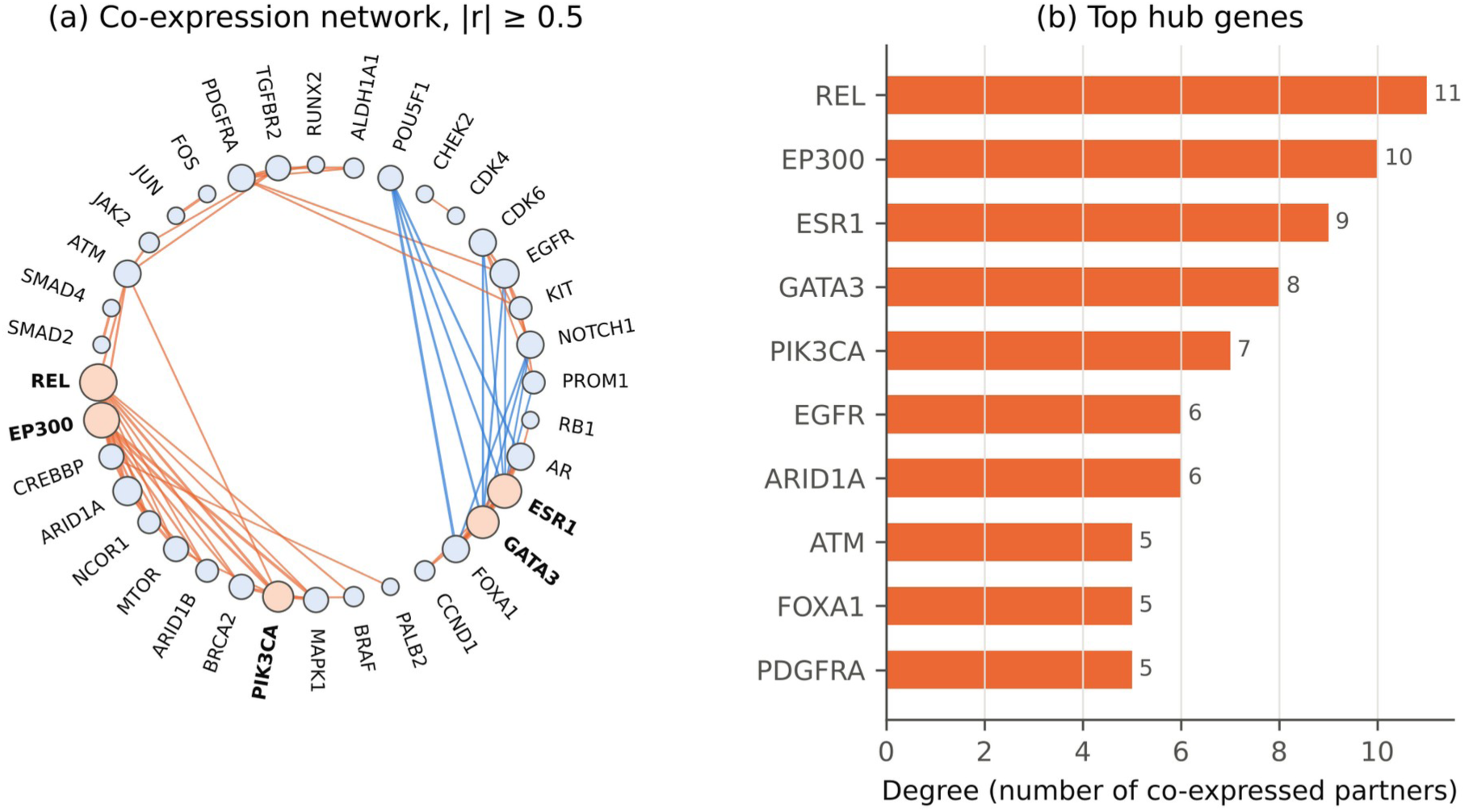
(a) Gene co-expression network at |r| ≥ 0.5 computed on the 860 training tumours; orange edges denote positive and blue edges negative correlations, node size is proportional to degree, and the five highest-degree genes are shaded orange. (b) Degree of the ten most connected genes; genes with equal degree are shown in arbitrary order. Edges represent co-expression, not regulatory interactions.

## Discussion

Using a leakage-aware pipeline on 1,075 TCGA-BRCA primary tumours, we found that a panel of 65 cancer-related genes has a modest ability to discriminate early-from late-stage breast cancer: the model with the highest test AUC, Logistic Regression, reached a ROC-AUC of 0.618 (95% CI 0.530–0.704) and a balanced accuracy of 0.578. Linear and non-linear models performed similarly, and none was significantly better than another, so these data provide no evidence that non-linear interactions among these genes add predictive information. These estimates are lower than the accuracies often reported for expression-based cancer-type or subtype classification [7–9], but they address a harder and different question.

Stage reflects the anatomical extent of disease—tumour size, nodal spread and metastasis—and also the time at which the tumour was detected. These properties are only partly encoded in the expression profile of the primary tumour, whereas molecular subtype and proliferation, which expression captures well, are related to but distinct from stage [3, 4]. A modest AUC is therefore an informative result rather than a failure: it quantifies how much stage information a cancer-gene panel carries under leakage-aware evaluation, and it indicates that any expression-based stage model is likely to require clinicopathological information to be useful. Our results also illustrate why accuracy is a poor guide in imbalanced problems [14, 15]: because 75% of tumours are early stage, a model that predicts early stage for every tumour attains 75% accuracy while having no discriminative value. Reporting ROC-AUC with confidence intervals, balanced accuracy, sensitivity, specificity and MCC, and fitting SMOTE and scaling only on training data [17, 18], gives a more honest picture of what the data support.

The three gene-level analyses answer different questions and should not be read as mutually confirming evidence. Permutation importance identifies genes that a particular fitted model relies on; differential expression tests whether a gene’s distribution differs between stage groups; and network degree measures how strongly a gene co-varies with other genes in the panel. In this study the three lists did not converge: none of the ten most important genes in the Gradient Boosting model was among the five highest-degree hubs, and no gene was significantly differentially expressed. This lack of convergence is itself consistent with a weak, distributed stage signal.

Several of the highlighted genes have known roles in cancer biology. *POU5F1* encodes OCT4, a pluripotency transcription factor linked to stem-like tumour cell states; *FLT3* encodes a receptor tyrosine kinase best known for its role in haematopoiesis and acute myeloid leukaemia; *NOTCH1* encodes a Notch receptor that regulates cell-fate decisions and is implicated in mammary tumorigenesis; *REL* encodes c-Rel, a subunit of the NF-κB transcription factor; *EP300* encodes a histone acetyltransferase and transcriptional co-activator; and *ESR1* encodes oestrogen receptor alpha, the defining marker of luminal breast cancer. These descriptions provide biological context only. The importance ranking comes from a model whose AUC confidence interval included 0.5 and is therefore exploratory; hub status reflects co-expression within a pre-selected panel and does not demonstrate regulatory activity; and any association with stage in this cohort is correlative. The data cannot show whether these genes drive progression or reflect differences in molecular subtype or tumour composition between stage groups, and none of them should be regarded as a validated stage biomarker.

The co-expression network should be interpreted with similar caution. Correlation-based networks describe co-expression, not regulation [31], and hub status in a network built from a pre-selected panel partly reflects how many related genes were included; for example, genes from the same pathway or lineage programme will tend to be highly connected because they were selected together.

### Limitations and Future Work

This study has several limitations. First, it uses a single cohort without external validation; independent cohorts are needed to confirm generalisability. Second, the gene panel was fixed in advance, which protects against selection leakage but limits discovery; whole-transcriptome feature selection nested inside cross-validation may capture additional signal. Third, the test set contained only 54 late-stage tumours, so performance estimates have wide confidence intervals, and the identity of the model with the highest test AUC may not be stable; cross-validation ranked SVM (RBF) slightly higher. Fourth, hyper-parameter tuning and the reported cross-validated AUC used the same folds, without nested cross-validation, and DeLong comparisons were not adjusted for multiplicity. Fifth, feature importance was assessed only for the Gradient Boosting model, not for the model with the highest test AUC. Sixth, TCGA samples were collected over many years under different editions of the AJCC staging system, and clinical covariates such as molecular subtype, grade and age were not included, so the univariate differential-expression results may be confounded by these factors. Finally, SMOTE creates synthetic samples by interpolation, which may not reflect real biological variation; the sensitivity analysis did not show a consistent advantage of SMOTE over class weighting.

Future work should validate the framework in independent cohorts, combine expression with clinicopathological variables and other omics layers, examine feature importance in the best-performing models, and test whether stage-associated genes add prognostic information beyond stage itself. These steps have not been performed in the present study.

## CONCLUSION

We present an interpretable, leakage-aware machine-learning framework for classifying breast cancer stage from gene expression. On 1,075 TCGA-BRCA primary tumours, the model with the highest test AUC (Logistic Regression) achieved a ROC-AUC of 0.618 (95% CI 0.530–0.704), and no classifier was significantly better than another. The study demonstrates that the expression of a 65-gene cancer panel in the primary tumour carries limited but measurable information about anatomical stage, and that imbalance-robust metrics are essential to describe such modest signals honestly. It does not demonstrate a clinically useful stage classifier, identify validated biomarkers or establish regulatory mechanisms: no gene was significantly differentially expressed after FDR correction, the feature-importance ranking is exploratory, and network hubs reflect co-expression only. External validation in independent cohorts and integration with clinicopathological variables are needed in future work before expression-based stage models can be considered for any clinical application.

## DECLARATIONS

### Data availability

The TCGA-BRCA data analysed in this study are publicly available from the UCSC Xena browser (https://xenabrowser.net), TCGA hub (https://tcga.xenahubs.net): gene expression dataset TCGA.BRCA.sampleMap/HiSeqV2_PANCAN and phenotype dataset TCGA.BRCA.sampleMap/BRCA_clinicalMatrix.

### Code availability

The analysis code, reference outputs and Supplementary Table S4 are openly available on GitHub at https://github.com/12345sri/breast-cancer-stage-ml-tcga and archived on Zenodo (release v1.0.0; https://doi.org/10.5281/zenodo.23456699) under the MIT licence. The notebook reproduces all reported tables and figures from the TCGA-BRCA input file described in the repository.

### Ethics statement

This study used only publicly available, de-identified data and did not require ethical approval.

## Acknowledgements

The results shown here are based upon data generated by the TCGA Research Network: https://www.cancer.gov/tcga.

## SUPPLEMENTARY MATERIAL

**Table S1:**
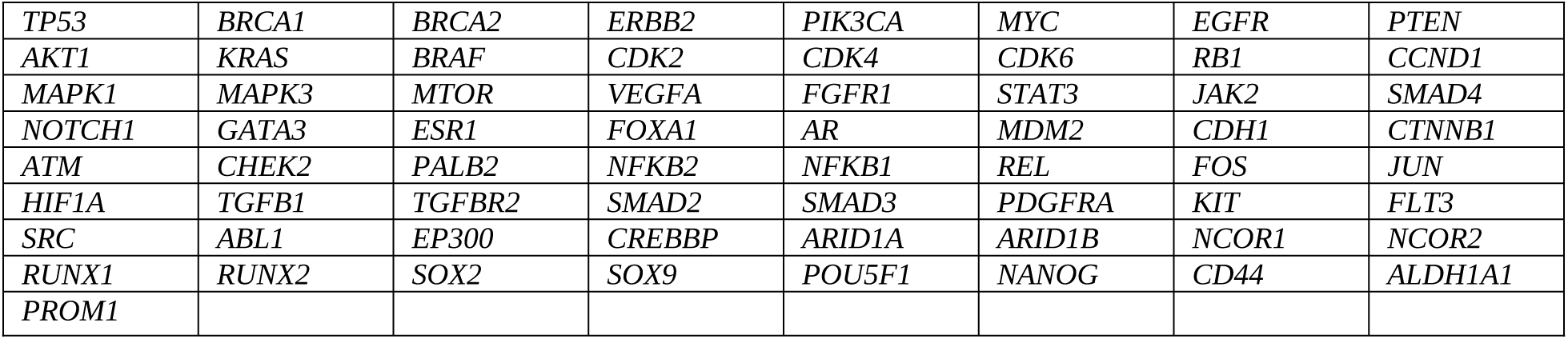
The 65 genes analysed (all genes were used as model inputs).

**Table S2:** Hyper-parameter grids searched with stratified five-fold cross-validation and the selected values. Fixed settings: Logistic Regression, L2 penalty; SVM, RBF kernel with probability estimates; Random Forest, 500 trees; Gradient Boosting, scikit-learn HistGradientBoostingClassifier without early stopping; random seed 42.

| Model | Grid | Selected |
| --- | --- | --- |
| Logistic Regression | C: [0.001, 0.01, 0.1, 1, 10] | C = 0.1 |
| SVM (RBF) | C: [0.1, 1, 10]; gamma: ['scale', 0.01, 0.001] | C = 10; gamma = 0.001 |
| Random Forest | max_depth: [None, 5, 10]; min_samples_leaf: [1, 5] | max_depth = None;<br>min_samples_leaf = 1 |
| Gradient Boosting | max_iter: [200, 400]; max_depth: [2, 3, 5]; learning_rate: [0.03, 0.1] | learning_rate = 0.1; max_depth = 5;<br>max_iter = 200 |

**Table S3:** Sensitivity analysis: test-set performance of class-weighted models trained without SMOTE, with thresholds selected as in the main analysis. Confidence intervals and DeLong tests were not computed for this analysis.

| Model | ROC-AUC | PR-AUC | Balanced accuracy | Sensitivity | Specificity | MCC |
| --- | --- | --- | --- | --- | --- | --- |
| Logistic Regression | 0.651 | 0.385 | 0.638 | 0.722 | 0.553 | 0.239 |
| SVM (RBF) | 0.625 | 0.378 | 0.607 | 0.630 | 0.584 | 0.186 |
| Random Forest | 0.615 | 0.369 | 0.570 | 0.537 | 0.602 | 0.122 |
| Gradient Boosting | 0.576 | 0.362 | 0.567 | 0.426 | 0.708 | 0.124 |

**Table S4** (full differential-expression results for all 65 genes) is provided as a spreadsheet file with this article.

