## Supplementary Table S4 for "An Interpretable Machine Learning Framework for Early- versus Late-Stage Classification of Breast Cancer from Gene Expression, with Co-expression Network Analysis"

**Table S4: Differential expression of all 65 panel genes between early-stage (AJCC Stage I–II; n = 805) and late-stage (Stage III–IV; n = 270) primary tumours (n = 1,075; two-sided Mann–Whitney U test), ordered by p-value.**

Means are on the log2 expression scale; Difference, mean late minus mean early; r, rank-biserial correlation (positive, higher in late stage); FDR, Benjamini–Hochberg adjusted p-value across the 65 genes. No gene reached FDR < 0.05. Values are rounded as in Table 3 of the main text (values that round to zero are shown as 0.00); unrounded values are provided in the accompanying spreadsheet.

| Rank | Gene | Mean early | Mean late | Difference | r | p | FDR |
|---|---|---|---|---|---|---|---|
| 1 | *CD44* | 0.52 | 0.28 | −0.24 | −0.14 | < 0.001 | 0.057 |
| 2 | *MAPK3* | 0.07 | 0.20 | +0.13 | +0.12 | 0.005 | 0.10 |
| 3 | *JAK2* | 0.37 | 0.21 | −0.16 | −0.11 | 0.005 | 0.10 |
| 4 | *TGFB1* | −0.40 | −0.27 | +0.13 | +0.11 | 0.007 | 0.11 |
| 5 | *PALB2* | 0.36 | 0.27 | −0.09 | −0.10 | 0.010 | 0.11 |
| 6 | *FLT3* | 2.13 | 1.74 | −0.38 | −0.10 | 0.014 | 0.11 |
| 7 | *ALDH1A1* | −1.89 | −1.65 | +0.24 | +0.10 | 0.014 | 0.11 |
| 8 | *AKT1* | 0.18 | 0.28 | +0.10 | +0.10 | 0.015 | 0.11 |
| 9 | *SOX9* | 0.73 | 0.54 | −0.20 | −0.10 | 0.017 | 0.11 |
| 10 | *SMAD2* | 0.23 | 0.15 | −0.08 | −0.10 | 0.017 | 0.11 |
| 11 | *TGFBR2* | −0.25 | −0.11 | +0.14 | +0.10 | 0.019 | 0.11 |
| 12 | *VEGFA* | −0.47 | −0.63 | −0.16 | −0.09 | 0.021 | 0.11 |
| 13 | *STAT3* | 0.32 | 0.24 | −0.08 | −0.09 | 0.024 | 0.12 |
| 14 | *FGFR1* | 0.69 | 0.85 | +0.16 | +0.09 | 0.029 | 0.13 |
| 15 | *SMAD4* | 0.01 | −0.05 | −0.06 | −0.09 | 0.031 | 0.13 |
| 16 | *PDGFRA* | 0.65 | 0.84 | +0.19 | +0.09 | 0.032 | 0.13 |
| 17 | *NFKB1* | 0.20 | 0.12 | −0.08 | −0.09 | 0.033 | 0.13 |
| 18 | *RUNX1* | 1.60 | 1.49 | −0.11 | −0.08 | 0.050 | 0.18 |
| 19 | *CHEK2* | 0.23 | 0.11 | −0.11 | −0.08 | 0.055 | 0.18 |
| 20 | *FOS* | −0.55 | −0.79 | −0.24 | −0.08 | 0.055 | 0.18 |
| 21 | *NFKB2* | −0.05 | −0.15 | −0.10 | −0.07 | 0.109 | 0.34 |
| 22 | *SOX2* | −3.21 | −2.85 | +0.36 | +0.06 | 0.126 | 0.37 |
| 23 | *BRCA2* | 0.76 | 0.60 | −0.16 | −0.06 | 0.138 | 0.39 |
| 24 | *MTOR* | 0.04 | −0.02 | −0.06 | −0.06 | 0.158 | 0.43 |
| 25 | *PIK3CA* | 0.35 | 0.27 | −0.08 | −0.06 | 0.165 | 0.43 |
| 26 | *PROM1* | 1.19 | 0.83 | −0.35 | −0.06 | 0.171 | 0.43 |
| 27 | *BRAF* | 0.26 | 0.21 | −0.05 | −0.05 | 0.196 | 0.47 |
| 28 | *GATA3* | 5.93 | 6.37 | +0.43 | +0.05 | 0.210 | 0.49 |
| 29 | *NANOG* | −0.06 | −0.12 | −0.06 | −0.05 | 0.217 | 0.49 |
| 30 | *SRC* | −0.38 | −0.44 | −0.06 | −0.05 | 0.232 | 0.50 |
| 31 | *ERBB2* | 1.38 | 1.53 | +0.15 | +0.05 | 0.239 | 0.50 |
| 32 | *ATM* | −0.04 | −0.12 | −0.08 | −0.05 | 0.254 | 0.50 |
| 33 | *CDK4* | −0.14 | −0.11 | +0.03 | +0.05 | 0.256 | 0.50 |

| Rank | Gene | Mean early | Mean late | Difference | r | p | FDR |
|---|---|---|---|---|---|---|---|
| 34 | *EGFR* | −1.97 | −2.19 | −0.22 | −0.04 | 0.272 | 0.51 |
| 35 | *REL* | 0.72 | 0.61 | −0.11 | −0.04 | 0.278 | 0.51 |
| 36 | *ARID1B* | 0.17 | 0.12 | −0.04 | −0.04 | 0.282 | 0.51 |
| 37 | *CREBBP* | 0.21 | 0.15 | −0.05 | −0.04 | 0.292 | 0.51 |
| 38 | *NOTCH1* | −0.65 | −0.65 | 0.00 | +0.04 | 0.308 | 0.52 |
| 39 | *SMAD3* | −0.43 | −0.35 | +0.07 | +0.04 | 0.312 | 0.52 |
| 40 | *KIT* | 0.62 | 0.49 | −0.13 | −0.04 | 0.343 | 0.56 |
| 41 | *MYC* | −0.03 | −0.10 | −0.07 | −0.04 | 0.359 | 0.57 |
| 42 | *CDK6* | −0.88 | −1.03 | −0.15 | −0.04 | 0.378 | 0.58 |
| 43 | *CTNNB1* | 0.21 | 0.16 | −0.05 | −0.04 | 0.383 | 0.58 |
| 44 | *POU5F1* | −1.66 | −1.89 | −0.23 | −0.03 | 0.400 | 0.59 |
| 45 | *NCOR2* | −0.25 | −0.24 | +0.01 | +0.03 | 0.443 | 0.64 |
| 46 | *RB1* | −0.17 | −0.10 | +0.07 | +0.03 | 0.508 | 0.72 |
| 47 | *KRAS* | 0.19 | 0.18 | −0.01 | +0.03 | 0.537 | 0.74 |
| 48 | *NCOR1* | 0.17 | 0.17 | 0.00 | +0.02 | 0.597 | 0.81 |
| 49 | *MDM2* | 0.18 | 0.18 | 0.00 | −0.02 | 0.609 | 0.81 |
| 50 | *TP53* | 0.21 | 0.21 | −0.01 | −0.02 | 0.625 | 0.81 |
| 51 | *ARID1A* | 0.25 | 0.24 | −0.01 | −0.02 | 0.689 | 0.87 |
| 52 | *ABL1* | −0.10 | −0.09 | 0.00 | +0.01 | 0.723 | 0.87 |
| 53 | *BRCA1* | 0.26 | 0.27 | +0.01 | −0.01 | 0.733 | 0.87 |
| 54 | *RUNX2* | 0.85 | 0.91 | +0.05 | +0.01 | 0.740 | 0.87 |
| 55 | *EP300* | 0.14 | 0.13 | −0.01 | +0.01 | 0.753 | 0.87 |
| 56 | *PTEN* | 0.24 | 0.22 | −0.01 | −0.01 | 0.762 | 0.87 |
| 57 | *FOXA1* | 4.93 | 5.49 | +0.56 | +0.01 | 0.775 | 0.87 |
| 58 | *JUN* | −0.31 | −0.32 | −0.01 | +0.01 | 0.776 | 0.87 |
| 59 | *AR* | 3.22 | 3.45 | +0.23 | +0.01 | 0.788 | 0.87 |
| 60 | *ESR1* | 5.16 | 5.60 | +0.44 | +0.01 | 0.837 | 0.90 |
| 61 | *CCND1* | 0.96 | 0.96 | 0.00 | −0.01 | 0.842 | 0.90 |
| 62 | *CDK2* | 0.17 | 0.15 | −0.02 | 0.00 | 0.905 | 0.93 |
| 63 | *HIF1A* | −0.71 | −0.70 | +0.01 | 0.00 | 0.909 | 0.93 |
| 64 | *CDH1* | 1.47 | 1.40 | −0.06 | 0.00 | 0.913 | 0.93 |
| 65 | *MAPK1* | −0.17 | −0.18 | −0.01 | 0.00 | 0.968 | 0.97 |
